# Novel diamond shuttle to deliver flexible bioelectronics with reduced tissue compression

**DOI:** 10.1101/435800

**Authors:** Kyounghwan Na, Zachariah J. Sperry, Jiaao Lu, Mihaly Vöröslakos, Saman S. Parizi, Tim M. Bruns, Euisik Yoon, John P. Seymour

**Affiliations:** Electrical Engineering & Computer Science Department, University of Michigan, Ann Arbor, USA; Biomedical Engineering Department, University of Michigan, Ann Arbor, USA; Biointerfaces Institute, University of Michigan, Ann Arbor, USA; The Neuroscience Institute, New York University, New York, NY, USA

## Abstract

The ability to deliver flexible biosensors through the toughest membranes of the central and peripheral nervous system is an important challenge in neuroscience and neural engineering. Bioelectronic devices implanted through dura mater and thick epineurium would ideally create minimal compression and acute damage as they reach the neurons of interest. We demonstrate that a three-dimensional diamond shuttle can be easily made with a vertical support to deliver ultra-compliant polymer microelectrodes (4.5 μm thick) in-vivo through dura mater and thick epineurium. The diamond shuttle has 54% less cross-sectional area than an equivalently stiff silicon shuttle, which we simulated will result in a 37% reduction in blood vessel damage. We also discovered that higher frequency oscillation of the shuttle (200 Hz) significantly reduced tissue compression regardless of the insertion speed, while slow speeds also independently reduced tissue compression. Insertion and recording performance are demonstrated in rat and feline models, but the large design space of these tools are suitable for research in a variety of animal models and nervous system targets.

## 1 Introduction

Implantable sensor arrays, especially when deployed in neural mapping studies, are most accurate and insightful when damage and disruption to nervous system circuitry is minimized. Minimizing damage and reactivity has been emphasized by recent global initiatives including the BRAIN Initiative, the NIH SPARC program, the GSK Innovation Challenge, and several DARPA funding programs, which have supported the development of new technology for mapping, monitoring, and/or controlling the nervous system with higher fidelity and longevity. Previous work demonstrated that size of an implant effects the neuron count and several glia markers of reactivity^1,2^. A precise implant size to tissue damage function is still missing, but evidence suggests that care should be taken to minimize the damage to blood vessels which densely permeate nervous tissue^3,4^. Central nervous system (CNS) neurons, for example, are located within ~15 μm of a capillary vessel^5–7^ so damage to the blood-brain barrier is unavoidable. When designing an implantable device, the penetrating structure must be stiff enough to reach the desired target with accuracy and, more challenging, piercing a tough fibrous outer membrane protecting the neurons. These challenges have opposing solutions – make the device as fine as possible for minimal bleeding and cellular disruption, yet ensure it is sufficiently stiff so as not to buckle or move off-target during penetration. The challenge is underscored when the covering membrane, the epineurium or dura mater in the peripheral or central nervous systems respectively, is left intact during insertion.

Epineurium and dura mater are made of collagen fibers interspersed with elastin and fibroblasts that are at once tough and flexible^8,9^. Cranial and spinal dura mater have a Young’s modulus in the range of 2-100 MPa^10^ (depending on species and location) and so compared to the parenchyma, whose modulus is ~5 kPa, provide the primary resistance to device insertion. In the feline dorsal root ganglia (DRG), the epineurial thickness is in the range of 20-100 micrometers (typically 80μm)^11^. The dura can often be separated with minimal damage to the parenchyma below with surgical training and skill, however the creation of dural flaps severs blood vessels extending into brain and allows the brain to swell and dehydrate ^4^. Furthermore, surgically removing the epineurium at the DRG or further into the periphery is not done in practice without damaging the nervous tissue. As such, a minimally invasive solution to penetrating through CNS and peripheral nervous system (PNS) membranes is needed.

State-of-the-art sensor arrays have taken different approaches to the problem of penetration when the membrane cannot be removed. One high-density and popular electrophysiology research tool is the “Utah” array^12,13^, which is a silicon array comprised of many ~70 μm diameter needles. It is inserted with a pneumatic insertion tool and can achieve an insertion velocity of 12 m/s^14^. The array has a wide base so it naturally stops when fully inserted but also may induce significant compression or impact trauma. No published quantification of the insertion force and compression distance exists, but significant bleeding has been reported^15,16^. Potentially related to this issue, the histology clearly shows encapsulation around the shank tips^15,17,18^ with a typical electrode-tip to axon distance of 40 to 150 μm. The resulting tissue restructuring around these microelectrodes prevents high-fidelity chronic recording, although it is still effective in recording population-based compound action potentials and for electrical stimulation^19^.

Another popular recording/stimulation array, but with a smaller footprint and greater electrode density, are planar silicon probes. Historically these have been called “Michigan” arrays but these micromachined silicon structures are now made by many groups^20–24^ and companies. They are typically inserted in the CNS using a manual manipulator at a moderate speed so as to monitor for buckling during insertion.

Given the risk of buckling or fracture, silicon planar probes are rarely used to penetrate tough membranes and there is even less known about the insertion mechanics. One study also demonstrated there is a chronic breach in the blood-brain-barrier for rigid intracortical structures ^25^.

New research has been focused on implanting flexible arrays with assistance from a variety of techniques. Given the impressive flexibility and stretchability of nerves, ganglia, and the spinal cord, researchers have attempted to insert flexible forms of microelectrodes. The most common insertion techniques are pulling the array through the tissue using thread and needle, a dissolvable stiffener, or a removable shuttle, otherwise known as a stylet. Several groups have demonstrated the pull-through technique for intrafascicular recording used a fine commercial needle and suture^26,27^. The diameter of these suture needles are hundreds of microns and risk causing significant trauma. As Wurth, et. al, carefully demonstrated in their histology the electrode to axon distance is typically 100 microns or more over time^27^. A dissolvable stiffener comes at the cost of greater volume given the temporary material is several orders of magnitude less stiff than many inorganics^28,29^, let alone the potential reactivity induced by the degraded polymer itself. The shuttle or stylet approach ^2, 30–33^ allows for stiffer materials and thus a smaller footprint than commercial needles or organic stiffeners. However, none of those approaches would allow for insertion through a tougher membrane without making the substrate much larger and at the cost a significantly more tissue damage.

### Our approach

This paper seeks to address the combined insertion challenge of minimizing the cross-sectional area and the tissue compression. This work adds to previous research using a shuttle approach but having a unique T-shaped geometry and material to form a three-dimensional ultrananocrystalline diamond (UNCD) shuttle with 13-fold greater stiffness than a similar cross-section made with conventional silicon (Fig. 1b). UNCD has the highest modulus of elasticity (800-1000 GPa)^34,35^ of any conformal thin-film and 5X and 3X higher than silicon^32,33^ and carbon^30^. We demonstrate a simple 2-mask method of fabrication (Fig. 1c). We also describe a biological model of blood vessel damage that others can use to predict damage from any arbitrary geometry. We also quantify the insertion dynamics in phantom and various tissues. Finally, we use the shuttle to demonstrate the delivery of and neural recordings from an ultra-compliant polyimide array (Fig. 1a). We expect this new ability to deliver higher-density bioelectronics with less compression and less vascular damage will improve the understanding of CNS and PNS function.

**Fig 1.**
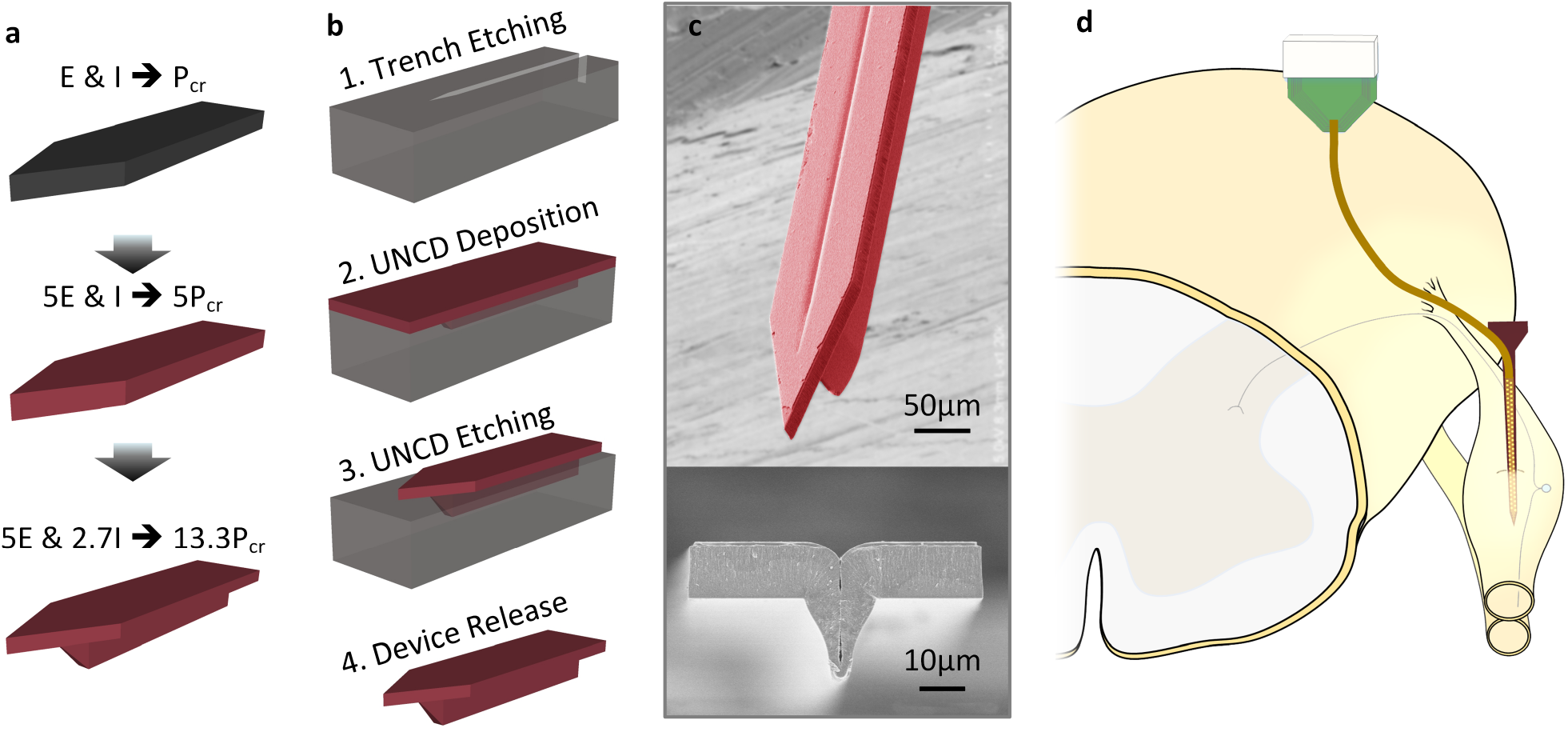
Insertion concept, design concept, and fabrication of a ultra-nano-crystalline diamond (UNCD) shuttle. **a,** Progression of design improvement from a simple silicon shuttle, with a buckling load of P_cr_, to an improvement of 13.3*P_cr_ by changing the Young’s Modulus, E, and moment of inertia, I. **b,** Process flow of UNCD shuttle. **c,** SEM of released UNCD shuttle. **d,** Insertion of flexible electrode array into a dorsal root ganglia (DRG) using a rigid shuttle and retraction.

## 2 Results

### Blood vessel damage model

While a reduction of area potentially leads to diminution of insertion damage to tissue, we have no biological evidence that this rule is still valid for T-shaped geometry. To address this, we modeled the damage using an available model of the density, size, and length of micro-vascular segments in the brain^6,7^. A three-dimensional micro-vascular network data for rat brain is publicly available.^36^ We overlaid the contour of a variety of devices onto the microvasculature maps provided initially by scanning electron microscope (SEM) imaging and 3D reconstruction. The model assumes that the device contour damages any vessel for which there is contact. This approach allows us to assess the performance of T-shape geometry in terms of a quantitative analysis of blood vessel invasion. The numbers of micro-vascular segments cut by the footprints of the shanks were counted throughout 1,000 insertion iterations with randomized angles and locations for two equivalently rigid geometries (Fig 2). The chosen T-beam cross-section contour was 11 × 65 μm on the planar portion and a vertical support that was 27.5μm-deep that tapered from 16 to 2μm-wide. This T-shaped UNCD (T-UNCD) was used for this model and throughout the study. The silicon device that has an equivalent buckling strength is 34-μm thick. All devices in this study are 65-μm wide which is generally the accepted width of a minimally damaging brain probe for chronic recording^23,37^. On average the simulation produced a 36.6% reduction in blood vessel damage caused by the T-UNCD relative to the Si-34 rectangular silicon device. Two important points of reference are a 15-μm thick silicon probe, which is sufficient for insertion in rodent models with the dura removed, and another commercially available neural probe that is 50-μm thick and intended for larger animal models. We compare these two against a T-UNCD structure that has equivalent stiffness with Si-15 (T-UNCD SMALL) and is considerably smaller in area. The simulation results were 1.4 vessels damaged compared to 5.8 vessels damaged by Si-15. By contrast, the 65×50-μm geometry caused damage that was 9.7-fold greater than the smaller T-UNCD (Supp. Fig. 2).

**Fig 2.**
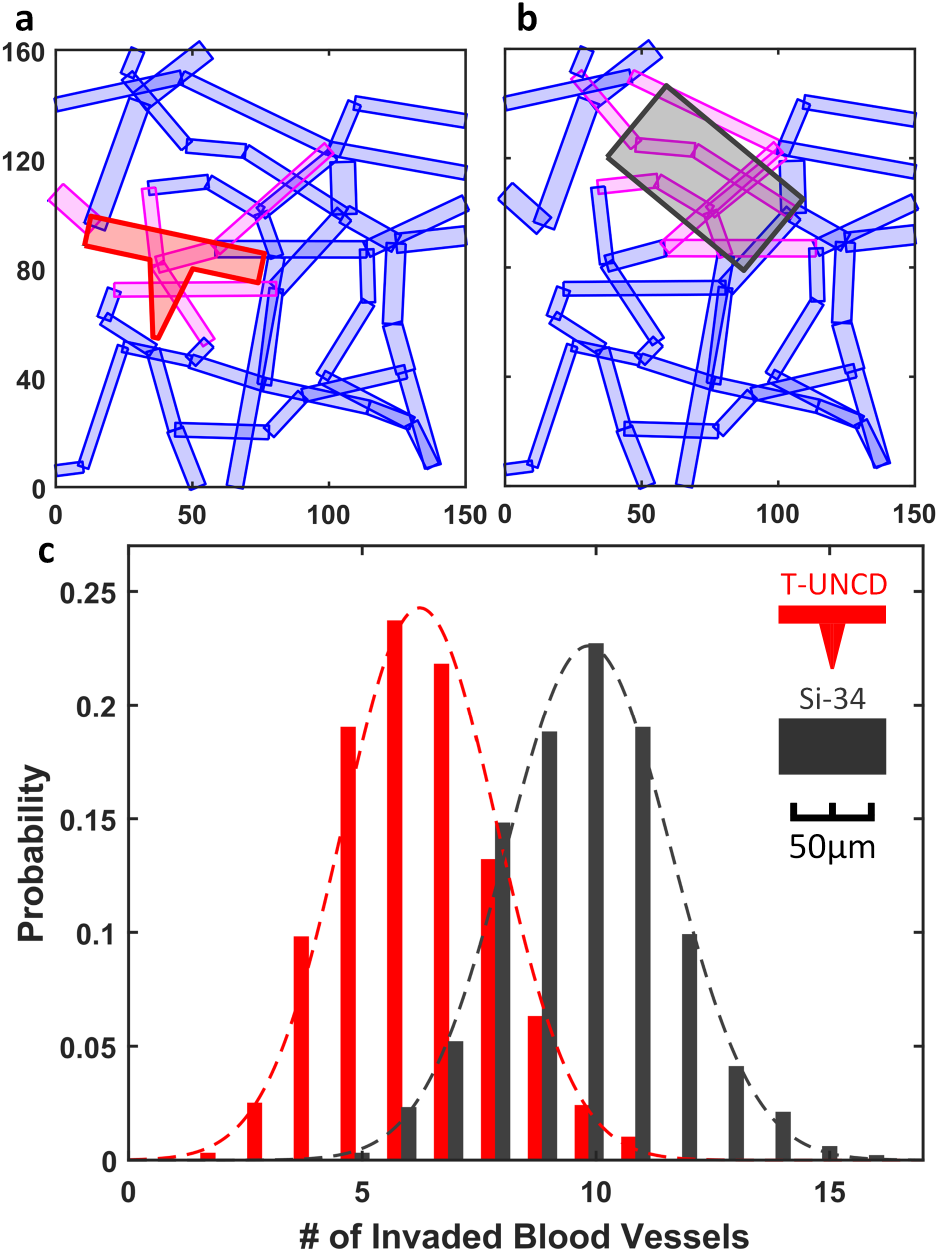
Simulation result of blood vessel invasion comparing Si (dark gray) and T-UNCD (red) having an identical buckling strength. Examples of invasion analysis with **a,** T shape with 65μm by 11μm on the planar portion and 27.5μm-deep, 16μm to 2μm-wide trapezoid for the vertical support, which has identical buckling strength with (b). **b,** 65μm by 34μm rectangle. **c,** Simulation result of 1,000 insertions yielded a Gaussian distribution of the number of invaded blood vessels. Mean values of two geometries from (a-b) were 6.23 and 9.84, respectively. Invaded blood vessels in (a-b) indicated in magenta. X and Y labels in (a-b) indicated the coordinates in μm.

### Creating 3-D UNCD structures

The diamond T-beam shape was successfully created by patterning a tapered trench into silicon and then filling it with a moderately conformal UNCD diamond. The conformality of UNCD deposition is related to the sticking coefficient and varies in a large range according to the deposition condition and the species involved^38^. Our specific conditions and sticking coefficient were unknown because the hot filament chemical vapor deposition method used for this work was provided by an outside vendor and the details were proprietary. We developed a method to model the deposition profile and estimate the sticking coefficient during deposition. This was critical for defining the proper silicon trench profile (Fig. 1c) and thus minimize void creation in the shuttle’s vertical support. Our deposition model (Supp. Note 1) fit best (R^2^>0.94) with a sticking coefficient, η_0_ = 0.079. The void formed was less than 0.4 μm across because of the trench’s taper angle of 70°. The angle of the silicon trench can be modified for deeper or wider geometries in the future by combining different doses of isotropic and anisotropic dry etching.

### Buckling load calculations & measurements

The most dominant mechanism of mechanical failure for insertion of a needle-shaped device is due to buckling. An insertion will end up with a failure when the buckling load is lower than the required insertion force. In this case, the shank keeps deflecting more without a load increase and breaks eventually when the deflection-induced stress exceeds the fracture strength of the material. Buckling loads for 15μm-thick planar Si (Si-15), 50μm-thick planar Si (Si-50), 11μm-thick planar UNCD (P-UNCD) and T-shaped UNCD (T-UNCD) shanks were measured (Fig. 3a). The characterization of buckling load was essential to analyze the results of insertion testing. Another goal of the buckling load measurement is the validation of the elastic modulus of UNCD and its structural design. The buckling loads of Si-15 and Si-50 were measured as −8.4% and +11.3% of the calculated value (Fig. 3a). The buckling load was calculated by equation (1) as:

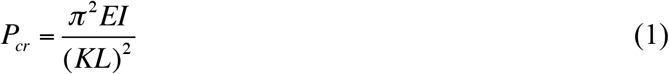

where E is Young’s modulus, I is the moment of inertia, K is the effective length factor, and L is the length. The K value is dependent on the boundary condition and it is assumed as a fixed-pinned condition in which K=0.699 but as discussed later is not always true. For consistency, all shuttles were also 1.5-mm long unless otherwise stated.

**Fig 3.**
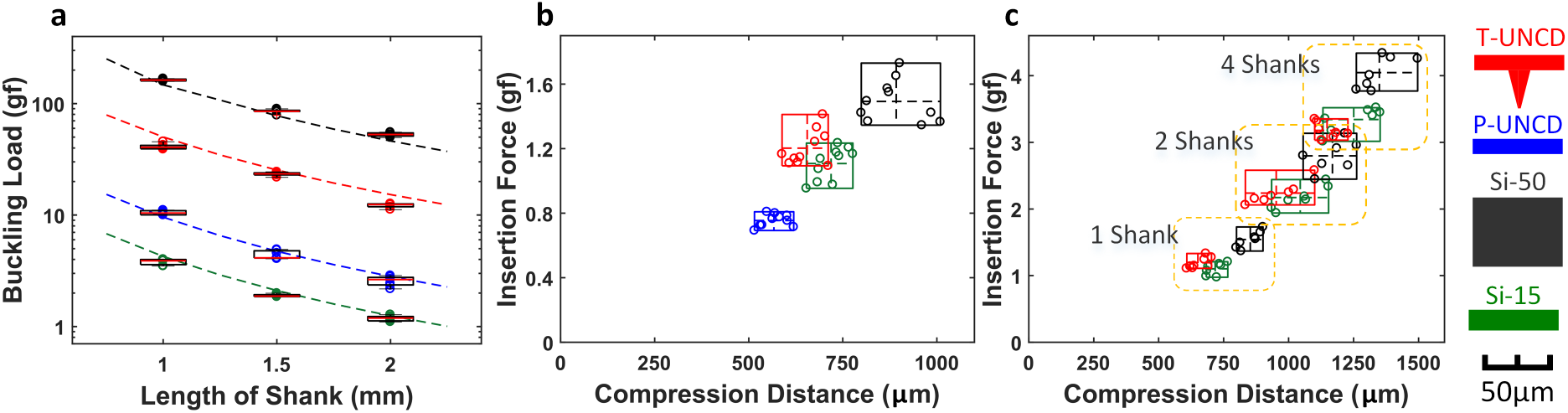
Mechanical characterization of various shuttles in a tissue phantom. **a,** Buckling load according to material, geometries and lengths, including two silicon geometries that are commercially available, N=7. **b,** Insertion force of single shank shuttles, N=10. **c,** Insertion force of 1, 2 and 4 shanks for three different probes, N=7 (planar diamond not tested). For 4-shanks, the force per shank was 0.8, 0.84, and 1.0 gf for T-UNCD, Si-15, and Si-50, respectively. The inter-shank spacing was 250 μm. Insertion speed was 0.01 mm/s in all cases.

The variation of thickness affects the buckling load significantly because the moment of inertia is proportional to the cubic of thickness. The range of UNCD’s Young’s modulus has been reported as 880-1000 GPa^34,39,40^ and our estimated value based on our known geometry best fit to 800 GPa. The measured buckling load of P-UNCD was 2.3% lower than that of the calculation, and it implies that the estimation for Young’s modulus is reasonable. The measurement of T-UNCD buckling load was 15.5% lower than the calculation and this is most likely due to the imperfection of the trench filling and the remaining void.

### Insertion force in phantom and feline DRG

*Ex vivo* tissue has a limited shelf life so many tests were conducted using a 100-μm thick PDMS sheet (Sylgard 184), to simulate dura and epinuerium, secured over a larger volume of agar gel. Our experiments showed that this phantom was stiffer than rat dura and almost as stiff as feline DRG epineurium (Table 1). Sylgard 184 (1-2 MPa)^41^ and agar have similar moduli as dura^10^ and parenchyma^42,43^. The chisel-shaped tips for all variations of shuttles had identical angles and the thickness varied based on film thickness, which was chosen because it is the standard tip of all Michigan-style probes.

**Table 1.** Average Peak Insertion Force ^A^

| Needle | T-UNCD |  |  |  | Pulled pipette<sup>B</sup> |  |
| --- | --- | --- | --- | --- | --- | --- |
| Tissue | Rat dura | Phantom | Cat DRG | Cat<sup>C</sup> dura | Rat dura | Phantom |
| Force, g | 0.68±0.22<br>N=6 | 1.2±0.1<br>N=10 | 1.9±0.7<br>N=6 | 1.4±0.2<br>N=2 | 0.40±0.03<br>N=2 | 1.7±0.2<br>N=8 |

Note A: Oscillation force data could not be collected
Note B: 2-μm diameter tip
Note C: Lumbo-sacral region of spinal cord

A typical force response is an increasing force during the initial insertion until an abrupt drop indicating a puncture event. This force-distance pattern of one or more punctures in soft-tissue is well-documented^44^. It was expected that the thinner tip required less insertion force than thicker geometries as was actually observed (Fig. 3b), though it was not linear. T-UNCD consistently showed multiple puncture events indicated by two force peaks. This corresponds to the tips of horizontal part and the vertical part of T-shape, respectively (Supp. Fig. 3b). The first peak force occurred at the range of P-UNCD insertion forces (0.75 g) and the second occurred following a further increase of force. The average peak insertion force for this T-UNCD was 1.2 gf and was 8.7% higher than Si-15 (Fig. 3b).

The insertion force of 1, 2 and 4 shanks for each cohort were compared to investigate the behavior of multi-shank insertion (Fig. 3c). Multiple shanks were 250 μm center-to-center from an adjacent shank. The force versus compression results, regardless of material and geometry, were less than a linear sum. Similarly, a three-dimensional assembly of six 4-shank UNCD arrays resulted in a decreasing force when normalized per shank (Supp. Fig. 4).

*Ex vivo* insertion tests with explanted feline L6, L7, and S1 DRG were conducted to evaluate the feasibility of epineurium penetration. The average insertion forces required for successful penetration of P-UNCD, Si-15, T-UNCD, and Si-50 were 1.15 ± 0.28, 0.96 ± 0.32, 1.92 ± 0.69 and 2.76 ± 0.61 gf, respectively (Fig. 4). However, Si-15 had the most failures (9 failures in 15 trials) followed by P-UNCD (4 failures in 19 trials), with no failures observed for the other two geometries. Our results indicate that T-UNCD will penetrate the feline epineurium with 100% success like Si-50 but with only about half of the cross-sectional area. P-UNCD and Si-15 will not reliably penetrate. In a qualitative *in vivo* test, the same T-UNCD design penetrated feline spinal cord (lumbar-sacral levels) nine times with zero failures where the dura is expected to be even thicker.

**Fig 4.**
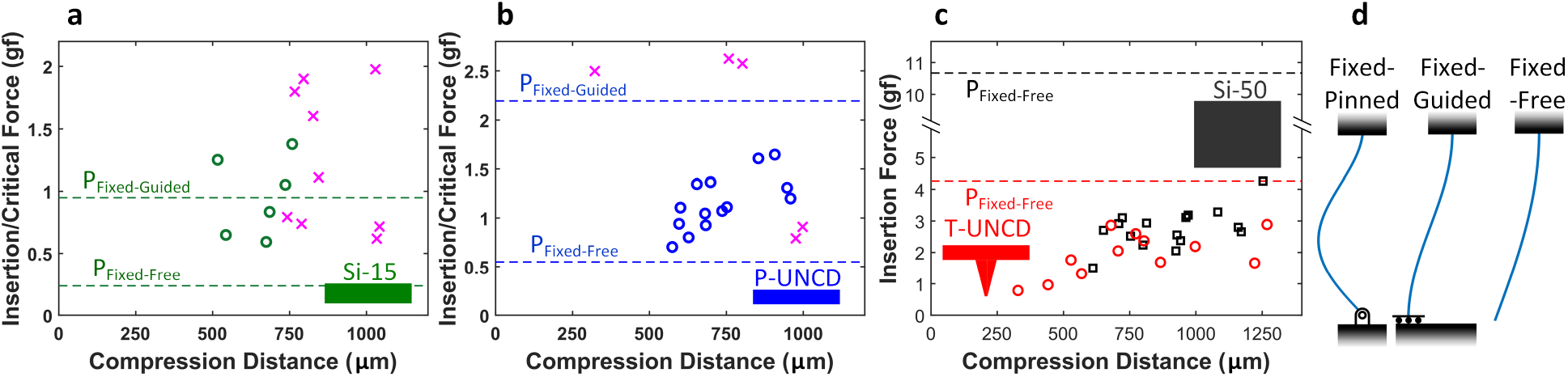
Result of ex-vivo insertions into cat DRG. **a,** 15μm-thick Si. “x” marks an insertion failure. **b,** 11μm-thick planar UNCD. **c,** T-shaped UNCD and 50μm-thick Si. **d,** Generic boundary conditions for the buckling load calculation (K=0.699, K=1, and K=2 respectively). The probe backend, at top, is fixed in each scenario and tissue is at the bottom. Theoretical limits noted as dashed lines in (a, b, c).

### Mechanical dynamics of insertion: speed and oscillation

In an effort to better understand the effect of linear speed and high-speed oscillation we modified our previous force sensor experiments. A piezoelectric motor capable of uniaxial movement with an amplitude of up to 70 μm was mounted onto a linear motor. A macro lens and video camera were also added to collect high-resolution compression information (Supp. Fig. 3c). Both linear speed and oscillation frequency were varied and the maximum compression distance (taken from annotated video images) was measured for each condition (N=6; Fig. 5). We found the greatest compression was 328 μm and occurred at 22 mm/s, linear motion only. At the other extreme, the compression was 124 μm at 0.01 mm/s and an oscillation of 200 Hz. This is a relative reduction of 62%. A graph of position and velocity over time for several tested motor parameters are provided (Supp. Fig. 6). A side-by-side video comparison of the UNCD shuttle at 0.01 mm/s with and without oscillation is also provided (Supp. Video 2).

**Fig 5.**
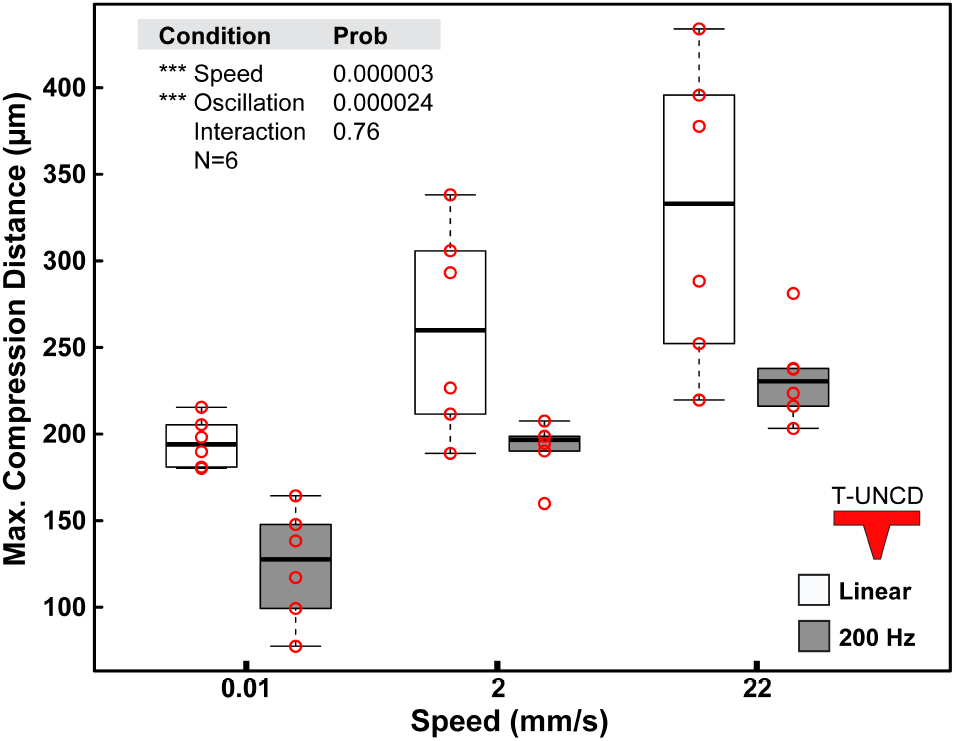
Result of T-UNCD phantom insertion with and without oscillation according to insertion speed. ANOVA results with an interaction term included. Both speed and oscillation were significant but the interaction term was not. Values for the maximum compression distance were taken from video analysis.

**Fig 6.**
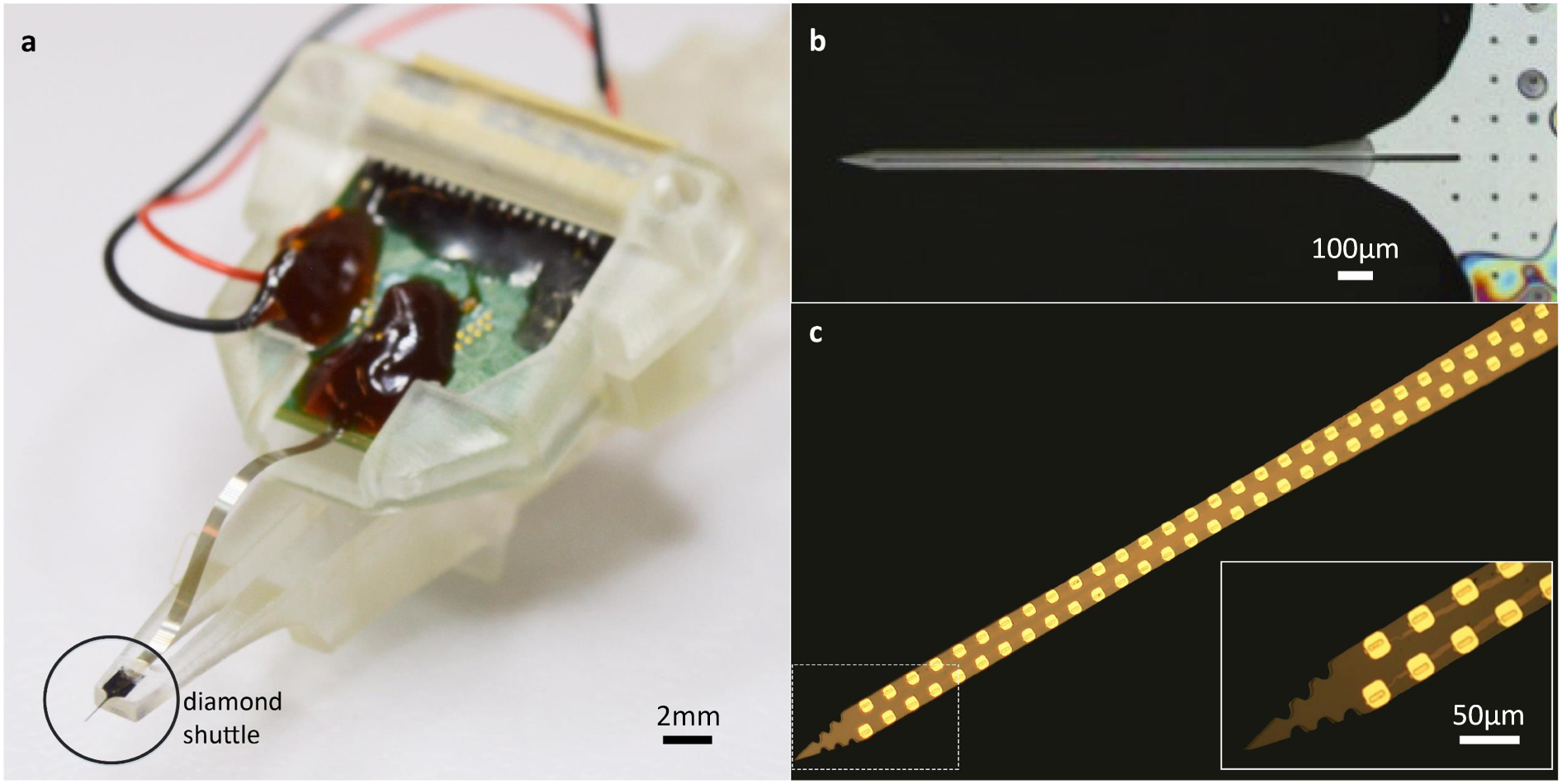
Assembly of high density flexible array mounted on UNCD shuttle. **a,** Photograph of entire assembly including the flexible polyimide array mounted on a UNCD shuttle. PCB is secured by a removable jacket that slides in a track of the shuttle jig. The insertion motor (not shown) is connected to the 3D printed shuttle jig. **b,** Top view of PEG-coated UNCD shuttle. **c,** Polyimide-based flexible array with 60 recording sites and (inset) zoomed view of the array tip.

Oscillation frequencies of 10 and 100 Hz were tested at 2 mm/s and a similar trend was observed with increasing frequency, which lowered the magnitude and variance of the compression (p=0.087, Supp. Fig 5). We also achieved a significant reduction of compression with a pipette (2-μm tip diameter) with 200 Hz oscillation (p=0.0007, Supp. Fig. 7).

### *In vivo* demonstration in feline DRG and spinal cord

The T-UNCD shuttle was inserted into the S1 or S2 DRG of three separate felines at 1.3 mm/s without oscillation. One animal had mechanical testing performed, which also included successful shuttle penetrations through the spinal cord dura (Table 1). In the other two animals, functional arrays were mounted on the T-UNCD shuttles (Fig. 6) and successfully recorded neural activity. Evoked activity was recorded from a variety of stimulus types across the span of the array, including cutaneous neurons from unique locations in the perineal region and base of the tail (Fig. 7). In the first experiment, a total of 28 different channels had a unit out of 42 functional channels, spanning 1 mm in depth across the array. We observed 7 putative units on subsets of these 28 channels, which were overlapping due to the closeness of the electrodes (40-μm pitch). In the second animal experiment, we observed approximately 40 putative units out of 48 channels spanning 1.06 mm in depth when the probe was in a fixed position. When the same probe was retracted, neural activity was observed on all 59 functional channels, including one putative bladder afferent (not shown). As described further in Supp. Note 3, in the second experiment we observed amplitudes greater than 1 mV and the median peak-to-peak amplitude for all experimental trials was 91 μV.

**Fig 7.**
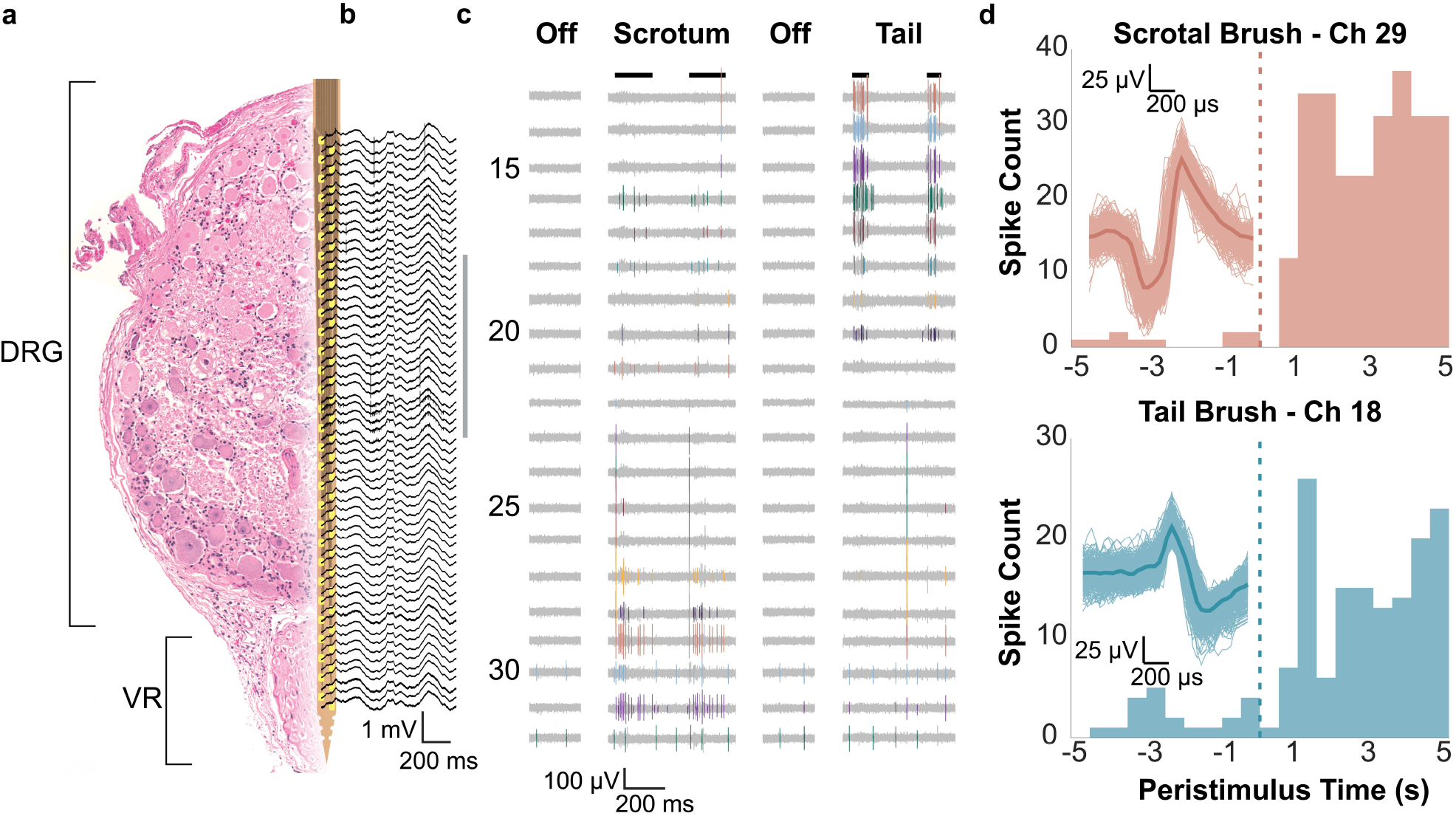
Recorded neural signals from feline sacral DRG for evoked stimuli. **a,** Diagram of the array in DRG. On the left is a representative hematoxylin and eosin stained cross-section of an example DRG with the ventral root (VR) below. Overlaid is a CAD diagram of the array, with array sites in yellow. The array and DRG are to scale. **b,** Wideband signal (highpass filter with 2 Hz cutoff) recorded during scrotum brushing. **c,** High pass filtered signals recorded during two sensory input trials (scrotum and tail brush strokes indicated by black lines at top). Channels shown are indicated by the gray vertical line to the right of (b). Manually sorted single units are in color. **d,** Peri-stimulus time histograms (PSTHs) for two channels and trial types, labeled and colored by channel numbers shown in (c). For both units shown, brushing starts at 0 s and continues through 5 s, including multiple individual strokes. Overlaid on each PSTH are the corresponding spike waveforms, with the mean waveform given as a dark thick line of the same color.

### *In vivo* demonstration in rat brain

We inserted the T-UNCD shuttle into the motor cortex through dura mater at 0.01 mm/s with and without oscillation (N=6) without fracture or failure. The depth of insertion was 1.5 mm. Without oscillation, the compression was 642 μm, and with 200 Hz oscillation the compression was 488 μm, a reduction of 24% (p=0.25, t-test). Separately, we also inserted the T-UNCD – polymer device through dura mater of anesthetized rats (2 insertions in each of 2 rats) also without failure. The flexible electrode was attached to the T-UNCD in the same way as in the feline surgery. After successful insertion the probe was separated from the UNCD shuttle (Supp. Video 3) and wideband (0.3 Hz – 3000 Hz) and single unit neuronal activity were recorded from the somatosensory cortex of the rat (N=1, Supp. Fig. 9a and b). The spontaneous neuronal activity was recorded for 30 minutes and 20 putative single units were found (Supp. Fig. 9).

## 3 Discussion

We demonstrated that in rat and feline models of dura and thick epineurium, our novel T-beam UNCD shuttle can deliver a thin, flexible high-density microelectrode array that would be especially useful in neural mapping studies. The reduced-footprint with high stiffness features of the shuttle exhibited better insertion features in tissue analogues and *ex vivo* neural tissue than standard probes.

There are also many design opportunities for thicker or thinner UNCD shuttles whose thickness, including the vertical support, can be in the range of 5 to 52 μm, which is potentially translated into the range of buckling load up to 102 gf. As we demonstrate in our results, important considerations like the boundary condition of a buckling model are especially important in the periphery. We also compare this shuttle approach to others.

### Buckling model highly dependent on boundary conditions

Throughout the *ex vivo* insertion tests with explanted feline DRG, many trials for Si-15 and P-UNCD resulted in failures (Fig. 4a, b) even though this was not expected based on the standard assumption of a probe tip “pinned” to stationary tissue (Fig. 4d). This commonly cited condition, where K = 0.699, may work for rodent brain implants but was incorrect for our *ex vivo* DRG experiments. Our results implied that the boundary condition at the contact between the tip and the tissue varied across trials presumably due to a varying angle of contact, the shape and curvature of the targeting tissue, and the friction of the tip at the surface. Also, that is a probable situation for *in vivo* insertion into peripheral nerve which will have similar variations. Adding to that risk is the breathing artifact observed in larger animal models. In these situations, the boundary condition is expected to be somewhere between fixed-guided (where the tip is pinned sufficiently to the tissue but the tissue itself rotates or moves;example failure in Supp. Video 1) or fixed-free (where the tip does not remain embedded in the surface and free to slip). This latter case is most likely due to a large insertion angle, which is challenging to avoid on highly curved surfaces. Also, a duller tip will not pin as effectively as a sharp one. The *ex vivo* insertion results suggested that it would be safe and wise to design the mechanics of a shuttle based on the worst-case boundary condition, especially when inserted into the PNS. This means the structure should be 8X stiffer than one designed for the fixed-pinned condition.

### Advantages of T-UNCD over other shuttle approaches

Mounting a polymer array on the shuttle and the separation after insertion is an important, practical issue for a successful *in vivo* experiment. Fortunately, a planar polymer array will self-align on a planar shuttle due to geometrical similarity and surface tension when dipped in a solution of water or polyethylene glycol. This may be a simpler approach than using a cylinder-shaped shuttle, e.g. tungsten wire or carbon wire^30,31^ where the fabrication is expensive and does not scale to multi-shank devices. Also the resulting cross-section is a rectangle tangent to a circle and thus creating more damage than either alone. T-UNCD has the advantages of both stiffness and a matching geometry for the high-density polymer arrays used here and by other groups ^26,27,33^. Also, compared to the stylet approach, the planar shuttle is more appropriate to deliver multi-shank electrodes readily enabling higher channel count. Furthermore the T-UNCD forms a sharp tip that is effectively notched to create a micropost. Luan et. al demonstrated the micropost but this version required focused ion beam etching which is slow and costly compared to large-scale microfabrication. Nonetheless the micropost concept is advantageous if the immediate release of a device is critical in an application and the potential of bending the polymer tip is not a concern. Future work may use the T-UNCD as a micropost in order to speed the release after insertion in feline DRG experiments where breathing movement was significant. In the above acute recording demonstration, the shuttle remained for almost an hour at a time and in each case the shuttle was broken from breathing movements. When the shuttle was removed within 5 minutes of implant, the shuttle was intact, but releasing the biosensor array within 5 minutes using polyethylene glycol was not possible in our experience. Thus the micropost approach with the UNCD would be ideal for chronic recordings in large animals with significant tissue movement.

We believe that T-UNCD is superior to other shuttles even when a large insertion force is not required (e.g., when dura mater is removed). Assuming 6mm length of a shank, buckling loads of T-UNCD SMALL and Si-15 are estimated as 0.19 gf and 0.16 gf, and the former reduces the blood vessel damage to 24% of the latter (Supp. Fig. 2). T-UNCD SMALL has 37% less cross-sectional area than 20μm-diameter tungsten wire which has an estimated buckling load of 0.18gf.

### Insertion dynamics with oscillation

The tissue compression in some scenarios was clearly visible, especially in feline *ex vivo* DRG samples where the small size of the ganglion underscored the surface movement (Fig. 4 and Supp. Video 1). Previous work in rats, albeit without an intact dura, demonstrated that low speed reduced force and tissue compression at a speed as low as 0.2 mm/s^45^. Therefore, our evident tissue compression at a much lower speed of 0.01 mm/s prompted further investigation beyond speed. The ability to reduce insertion force and compression with oscillations has been documented in mosquitos^46^ and demonstrated with a small needle attached to a vibratory actuator with a kilohertz frequency^47^. In the former case, slow oscillations of up to 15 Hz using a three-part cutting system make a fascinating but complicated micromachine. In the latter case, the oscillations were 600 μm in amplitude and lateral to the direction of insertion. While this reduced both compression and the insertion force, this risks damage to the nervous tissue. To address this, we employed a piezo stage with motion only in the direction of penetration and tested for the first time the combination of speed and frequency.

To study this required an accurate measure of compression. Initially, the experiments only collected force versus position data with low resolution video recording. In those experiments we could not measure the maximum compression distance but only the total distance the probe moved before the peak force was observed. This over-estimates the compression distance since slow or small rupture events occurred before a larger rupture event, as others have reported^45,48^. Later we added video recording and a stabilized 4X macro lens (Supp. Fig. 3c). From this the maximum compression (Fig. 5 and Supp. Fig. 5) could be more accurately measured with annotated video images and on-screen measurements.

For some time, some neuroscientists have used slow-speed insertion for its perceived reduction in dimpling and tissue strain^49^. Our data confirms that slow speed, independent from oscillations, reduced the compression distance even in the context of a tougher membrane. Interestingly, high-frequency oscillation creates higher speed but in this case also significantly reduced compression and variability. The kinematics of the movement is straightforward to model (Supp. Fig. 6) and the maximum instantaneous speed and acceleration is calculated in Supp. Table 1. Given that the components of the insertion force includes the compression force (non-linear spring model), friction, and cutting forces^44^, a more careful study should be conducted to isolate the effect of speed and oscillation on each. Further research should also measure the acute damage to the blood-brain-barrier^4,25,50^ over this range of motion. Until more is known about the effect of oscillation on the vascular system, the best strategy would be to simply turn off oscillation after the initial rupture event is detected.

## 4 Outlook

High-density bioelectronics for the nervous system have a long way to progress before we achieve biomimetic scale, equivalent tissue mechanics, and effectively no tissue-reactivity but the modeling and materials work shown here is an important step in that direction. The results presented here demonstrate the highest-density and most flexible polymer array to date—60 traces and 65-μm wide on a substrate only 4.5-μm thick—inserted through the toughest nervous system tissues to obtain neural signals. We have developed a versatile T-shaped ultrananocrystalline diamond shuttle which provides a 56% reduction in cross-sectional area compared to a planar silicon shuttle with equivalent insertion strength, and still stiff enough to insert into different neural structures in small and medium animal models with the dura and epineurium intact. This same structure will theoretically reduce micro-vascular damage by 37% compared to standard probes. We also quantified the benefits of slow insertion and demonstrated for the first time the use of uni-directional oscillation. This oscillation significantly reduced phantom tissue compression by 31%, showing the potential for limiting neural tissue impact for *in vivo* use. More broadly, the use and deployment of advanced bioelectronics as developed here is critical for high-resolution scientific research into the nervous system and will contribute to future clinical devices in the treatment of neurological damage and disorders.

## 5 Methods

### Blood-vessel damage model

A publicly available three-dimensional micro-vascular network data for rat brain^6,7,36^ was used to simulate the blood vessel invasion induced by the insertions of various geometries. For simplicity, the 3-D model was projected into a 2-D plane assuming that the insertion was vertical and straight. 1,000 insertion simulations were conducted per geometry and the inserted geometry was randomly located in terms of x, y locations and rotation per trial. A segment of blood vessel was classified as damaged when a device crossed the entire width of a segment. The distribution of the number of damaged blood vessels was fit to a Gaussian distribution to extract the mean value and the deviation.

### UNCD fabrication

The process flow is illustrated in Fig. 1c. First, standard photolithography defined a trench pattern on a 4” silicon wafer. A tapered trench was formed on silicon wafer by the combination of DRIE and isotropic dry etching. Smooth sidewalls were achieved by thermal oxidation and removal followed by another thermal oxidation for sacrificial layer growth. Afterward, 11-μm thick UNCD film was deposited by hot filament chemical vapor deposition (CVD) at Advanced Diamond Technologies, Inc. (Romeoville, IL). Then, 2 μm-thick plasma-enhanced CVD silicon dioxide was deposited at 350°C to be used as a hard mask for UNCD etching. Following photolithography, silicon dioxide and UNCD were dry-etched sequentially using C_4_F_8_ and O_2_ chemistries, respectively. The UNCD-only devices were released in buffered hydrofluoric acid by soaking for 6 hours and carefully rinsed with deionized water. A deposition model of UNCD in a trench is described in Supp. Note 1.

### Mechanical characterization

The inital setup for mechanical characterization was built using a precision load cell (Model 31, Honeywell, 50 g maximum), a digital multimeter (34011A, Agilent) and a motorized translation stage (MTS50-Z8, Thorlabs). This setup was used for data collected in Fig. 3, 4, and Supp. Fig. 4 and illustrated in Supp. Fig 3a. For the oscillation and speed studies (Fig. 5, Supp. Fig. 5 and Supp. Fig. 7) we used a piezostage HS1.70 from Piezoconcept (Lyon, France) mounted on the linear motor. The piezomotor was driven by a function generator where a 10V sine input produced a 70-μm PP amplitude. The silicon or UNCD shuttle was fixed to the end of load cell and the voltage output of the load cell was recorded through the ADC (225 samples per sec.) while the stage transfers the counterpart. The exerted force and travel distance were calculated by conversion of recorded data. The velocity and acceleration of the linear motor were 0.01 mm/s and 0.1 mm/s^2^ respectively.

A tissue phantom was formed by a 100μm-thick Sylgard 184 sheet secured over 0.7% agar gel. The *ex vivo* DRG and spinal cord tissues were used within 72 hours of extraction from felines used for other research purposes. All samples were kept in saline and continually rehydrated during an experiment to minimize mechanical degeneration or stiffening of tissues.

The recorded videos and the force-distance plots were used for the failure criteria of *ex vivo* DRG insertion. An insertion trial was classified as a failure when buckling was obvious from the video. In such cases, it was also evident in the force plot as there was no rupture event, i.e. drop in force over time. In other cases the shuttle fractured due to large bending.

### Maximum compression distance measurement

To capture more precise compression data during oscillation we used a Canon ES Rebel 5 with a macro lens (MP-E Macro 65mm 1:2.8) set to 4X zoom mounted on an optical bench (Supp. Fig. 3c). The insertion process was recorded over the full length of penetration, 1.5 mm. These videos were then analyzed in Photoshop. Initially a line marking the point of contact was annotated over the video, then a second line was added at the point of maximum compression. A pixel to pixel measurement was made in Photoshop and converted to distance. The pixel to distance ratio used in these videos was identical for all and determined by the known width of the shuttle (65 μm). The angle between the camera lens and phantom surface was measured (within 1 degree) and used to correct for the final maximum compression value; the average angle was 7 degrees.

### Polymer array fabrication

A flexible electrode array was fabricated broadly as we and others have described^31,51^. One layer of 2.2 μm-thick polyimide (PI-2610 from HD Microsystems) was spin-coated on a sacrificial layer and fully cured at 350°C. This thin, biostable PI is BPDA-PDA (3,3′,4,4′-biphenyltetracarboxilic dianhydride with p-phenylene diamine)^52^. This variety of polyimide was chosen because of its high yield strength, flexibility, and low water absorption^53^. Metal patterning was formed in several steps depending on resolution requirements. 500-nm lines at 65-nm thickness were formed using a GCA stepper i-line lithography tool. This high resolution enabled 60 traces at a maximum width of 70 μm. Future work could also employ multi-layer metal. Wider and thicker metal was patterned using a Seuss MA-6 contact aligner over the interconnects and bond pads. A TiO_2_ adhesion layer above and below metal traces was used to presumably improve adhesion, although not yet validated. The top PI-2010 layer was also 2.5-μm thick. The PI was etched by oxygen plasma to define the shape and open the electrical contact sites. Ti/Au electrode metal was patterned by a liftoff process and devices were released by the removal of the sacrificial layer. The electrode pitch was 40 μm. In order to decrease the impedance of the recording electrode sites, PEDOT-pTS was coated on Au electrodes as described elsewhere^54^. The impedance in PBS was lowered to 24.0 ±1.45 kΩ from 1.98±0.752 MΩ of Au electrode (Supp. Fig. 8).

### Mounting polymer array on UNCD shuttle

The ultra-flexible electrode array was electrically assembled on a PCB and then mechanically assembled on a UNCD shuttle for *in vivo* insertion and recording. First, the PI arrays were ball bonded onto a printed circuit board^55^ and the assembly was placed into a 3D-printed jacket designed to slide in the track of a shuttle jig (Fig. 6a). Separately, a UNCD shuttle was aligned and mounted on a 3D-printed shuttle jig followed by dip coating into a beaker of PEG (12,000 MW) heated to 70-90°C (Fig. 6b). The PI array assembly was roughly aligned over the shuttle, aligning the vertical position by sliding the array assembly in its track on the jig. This position was then fixed to the shuttle backend by a small drop of PEG (4,000 MW). Precise self-alignment was achieved by once again dip-coating in heated PEG (12,000 MW). The resulting film was again 1 to 2 μm thick and transparent. This self-alignment achieved an accuracy of at least 5-μm.

### *In vivo* experiment, feline DRG

Procedures were performed in three adult, intact felines (male, 4.6-6.1 kg). Neural recordings were performed in two animals while only insertion testing was performed in a third. All steps were approved by the University of Michigan Institutional Animal Care and Use Committee (IACUC; protocol PRO00007296). As described elsewhere^51^, felines were sedated with a ketamine-butorphanol-dexmedetomidine intramuscular dose prior to intubation and maintenance under isoflurane anesthesia (2-4%) with an intravenous saline drip (5-10 mL/kg/hr) and vitals monitored and maintained within normal ranges. A laminectomy was performed to expose the lumbosacral (~L7-S3) DRG. Bone screws were inserted into the iliac crests bilaterally. The feline was placed into a custom testing stand^56^ which supported the chest. Stainless steel wire was wrapped around each iliac crest bone screw and fixed to the stand, allowing for the lower abdomen and hind limbs to be suspended. This positioning reduced the relative motion due to breathing that would occur at the DRG. A jig holding a shuttle-mounted array was attached to a linear inserter (M-230.25, Physik Instrumente, Karlsruhe, Germany) mounted on a micromanipulator (KITE-R, World Precision Instruments, Sarasota, FL, USA), which was used to position the array just above a DRG (S1 or S2). The inserter was driven at a rate of 1.3 mm/s to insert the array and shuttle to a depth of 1.4 mm through the DRG. A variety of sensory stimuli were applied to activate sensory neurons, including brushing of different cutaneous dermatomes related to the DRG implanted (scrotum, anus, base of tail), infusion of saline into the bladder, and skin-surface electrical stimulation. Neural signals were sampled at 30 kHz with a Neural Interface Processor (Grapevine, Ripple, Salt Lake City, Utah, USA). After an experiment neural signals were first sorted in Offline Sorter (Plexon, Dallas, TX, USA) prior to analysis with custom scripts in MATLAB (Mathworks, Nantick, MA, USA).

### *In vivo* experiment, rat dura

The experiment was approved by the Institutional Animal Care and Use Committee at the University of Michigan (protocol number: PRO00007275). Two male wistar rats (600 g) were implanted with a UNCD-polyimide electrode under isoflurane anesthesia (1.2 – 1.5%). No prior experimentation had been performed on this rat. Atropine (0.05 mg/kg, s.c.) was administered after anesthesia induction to reduce saliva production, and the rectal temperature was monitored and kept constant at 36–37 °C with a DC temperature controller (TCAT-LV; Physitemp, Clifton, NJ, USA). Proper depth of anesthesia was maintained by confirming the lack of nociceptive reflex. A craniotomy was drilled (4 by 4 mm), carefully paying attention to not damage the dura mater. The hole was cleaned and filled with sterile saline solution (0.9%, MWI, USA) and kept wet until the custom-electrode was implanted at 1 mm posterior from bregma and 2.8 mm lateral of the midline, in the S1FL area of the somatosensory cortex.

The electrode was inserted to a 2 mm depth in the cortex using the custom motor system (0.01 m/s, 200 Hz oscillation, Supp. Fig. 3c). The insertion/dura penetration process was video recorded (Canon ES Rebel 5 camera was attached to a Zeiss OPMI-1 microscope, 31.25x total magnification). In one of the rats, the separation process was initiated by applying sterile saline on the array ten minutes after insertion. Once the upper part of the polyimide was separated the UNCD shuttle was slowly moved (50 μm up and down) until a complete separation was visible under microscope. Then, the shuttle was retracted while the recording electrode was kept in place (Supp. Video 3). The recorded signals (n = 60 channels) were amplified (200X gain) and stored after digitization at 20 kHz sampling rate per channel (RHD2132 and RHD2000 Evaluation System, Intan Technologies, Los Angeles, CA).

The recorded data were analyzed by custom-written scripts in MATLAB (MathWorks, USA). Neuronal spikes were detected from the digitally high-pass filtered signals (0.5–3 kHz) by Spikedetekt2 (https://github.com/klusta-team/spikedetekt2). Detected spikes were automatically sorted using KlustaKwik2^57^, followed by manual adjustment of the clusters using KlustaViewa software^58^ to get well-isolated single units (multi-unit and noise clusters were discarded).

## Acknowledgments

The authors thank the staff of the Lurie Nanofabrication Facility for their tool support and technical advice when needed. We thank Paras Patel for his help in PEDOT-pTS coating. We also thank Ahmad Jiman, Zhonghua Aileen Ouyang, and Lauren Zimmerman for their help in 3D printing materials for and/or data collection in feline experiments. This work was supported in part by the National Institute of Biomedical Imaging and Bioengineering of the National Institutes of Health (R21EB020811, and SPARC program Awards U18EB021760, OT2OD024907, and OT2OD023873), and Seed Funding for Innovative Projects in Neuroscience from the University of Michigan Brain Initiative Working Group (MiBrain). The content is solely the responsibility of the authors and does not necessarily represent the official views of the National Institutes of Health or the University of Michigan.

## Supplemental Information

### Supplemental Note 1: Mounting polymer arrays to the shuttle

Regarding the temporary adhesion of a polymer array onto the diamond shuttle, we describe our PEG mounting process below. The dissolution rate of PEG has to be carefully controlled so that the array doesn’t separate prematurely during insertion and the dissolution and separation happens within a reasonable time after insertion. For cat DRG, this is not trivial because there are several hundred microns of relative DRG position shifting as the vertebral region is coupled to the body’s movement during breathing. This movement is enough to either cause damage to the tissue or fracture the shuttle. We found that if the UNCD shuttle is removed within five minutes of insertion it did not break in three of three attempts. When left for longer, the shuttle fractured. Comparatively, a head-fixed rat surgery had no time limit and movement was hardly visible(Supplementary Video 3), although Gilletti et. al found this to be around 10-30 μm and varied with anesthesia^1^. We tested the molecular weight of PEG (4,000, 8,000, and 12,000 MW) and found the best result for our purposes to be dip coating with 12,000 MW. For rapid removal, using the T-UNCD as a micropost is expected to be as effective as other microposts^2,3^ except it would also be stiffer and/or smaller.

**Supplementary Figure 1.**
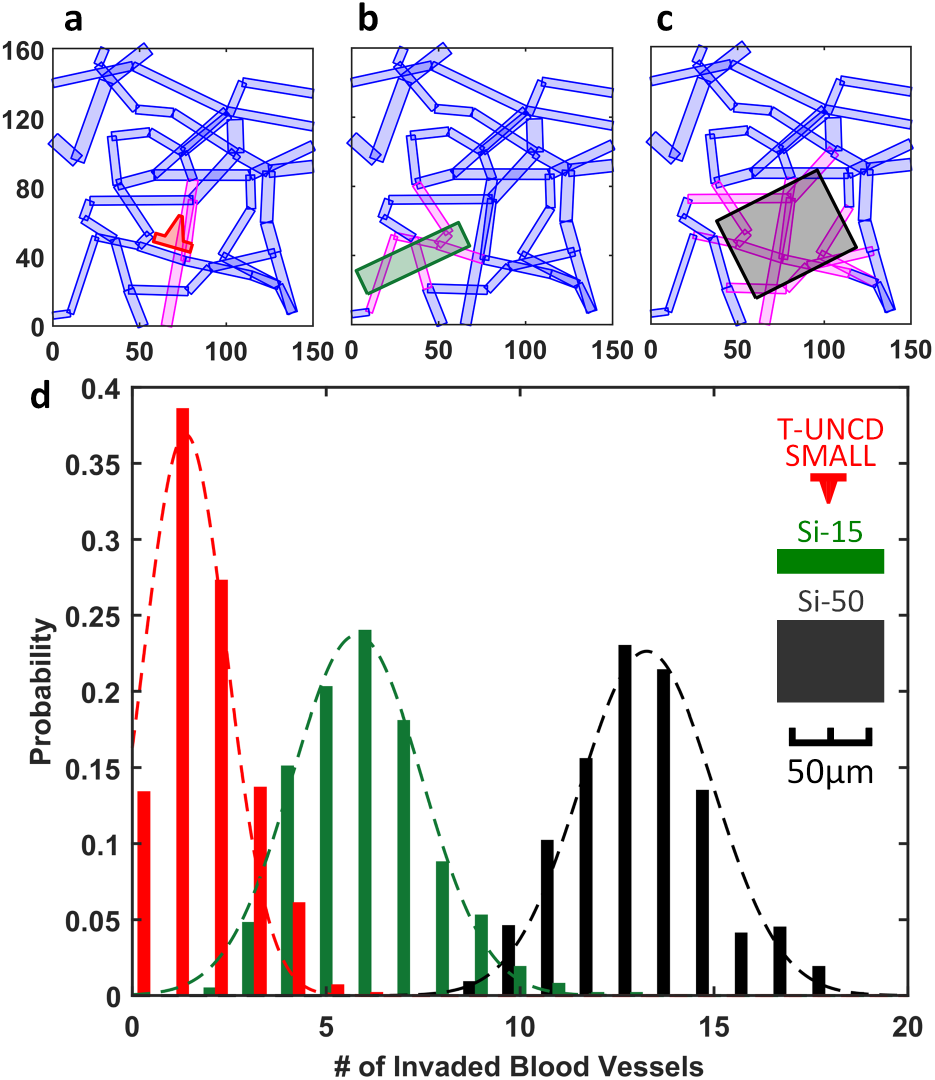
Simulation of blood vessel invasion based on various probe cross-sections. Examples of invasion analysis with **a,** T shape with 22 × 5 μm on the planar portion and 13.5μm-deep, 11 to 2μm-wide trapezoid for the vertical support (red, T-UNCD SMALL). **b,** 65μm by 15μm rectangle (green, Si-15). **c,** 65μm by 50μm rectangle(black, Si-50). **d,** Simulation of 1,000 insertions result in a Gaussian distribution of the numbers of invaded blood vessels. Mean values of three geometries from (a-c) were 1.4, 5.8, and 13.3, respectively. Invaded blood vessels in (a-c) indicated in magenta. X and Y labels in (a-c) indicated the coordinates in μm.

### Supplemental Note 2: Limits to conformal trench filling with UNCD

The sticking coefficient (η_0_) is a property of CVD deposition and is the most important parameter in determining a film’s conformation around 3-D structures. η_0_ is dependent on pressure, temperature, and the nature of each species, etc. Theoretically, the shape of the trench and the sticking coefficient determine the deposition profile. As mentioned in the Results section, the sticking coefficient of UNCD deposition varies in a large range and was unknown for the system we used at the time of deposition. The challenge of trench filling of UNCD is the moderate conformality of the UNCD film. Our model estimated that γ= 0.079, which is a moderate sticking coefficient^4^. We resolved the issue by creating a tapered trench (lower aspect ratio) but the curved shape of the T-beam also minimizes high-stress points that would exist around a sharp T-beam geometry. Furthermore, characterizing the critical factors contributing to the sticking coefficient is a long experimental process best done when the process is done in-house as opposed to a commercial service.

Ganguli et al^5^ and Komiyama et al^6^ suggested the governing equation of deposition model based on the mass conservation and it was reformulated as shown in (2). It was assumed that the width of trench is constant along with the depth as well as time. The assumptions are inappropriate due to tapered profile and increasing film thickness over time. The derivative equation becomes unsolvable form and analytical solution cannot be derived.

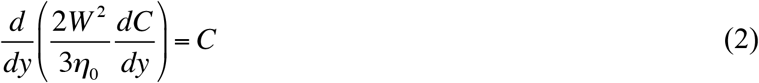

where W is the width of trench, η_0_ is the sticking coeffieient, and C is the concentration of reactant.

Instead, we converted the derivative equation to a discrete form and iterated the calculation according to the depth and the time. The film thickness deposited on discrete coordinates of y during a unit time was calculated with the fixed width and then the width was updated considering the deposited film for the following calculation during the next unit time. The calculation was iterated until the time when the targeted total thickness was achieved.

The model predicted a slight hollow in the middle of the trench since the etched profile was not linear and the sidewall angle of some portion was above the critical angle (70̌) for void-free filling even though the averaged angle was more tapered. Since a small void has a small change in the moment of inertia, we also expect a small loss of mechanical strength.

Finally, it is noted that the residual stress of UNCD film is not consistent during deposition and the resulting gradient make it challenging to control or compensate for residual stress with a thick film UNCD. The moderate sticking coefficient and stress gradient will eventually limit the aspect ratio and film thickness so the upper limit of strength we estimate to be in the range of 65 gf based on our current film properties. This is sufficient for all of the animal models for which we currently have data for.

**Supplementary Figure 2.**
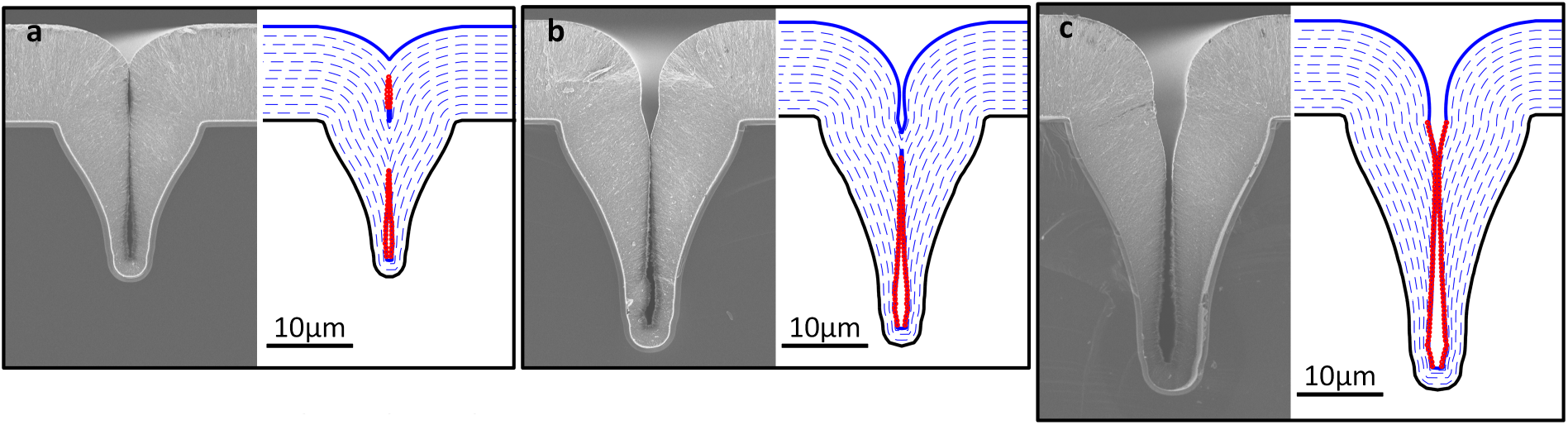
Simulation result of UNCD profile modeling. **a,** design A with 18.3μm-deep trench, R^2^=0.957. **b,** design B with 26.9μm-deep trench, R^2^=0.946. **c,** design C with 32.2μm-deep trench, R^2^=0.943. Blue solid line is the final outline of UNCD film. Blue dashed lines show the progression of deposition. Red lines indicate the measured profiles of UNCD film.

**Supplementary Figure 3.**
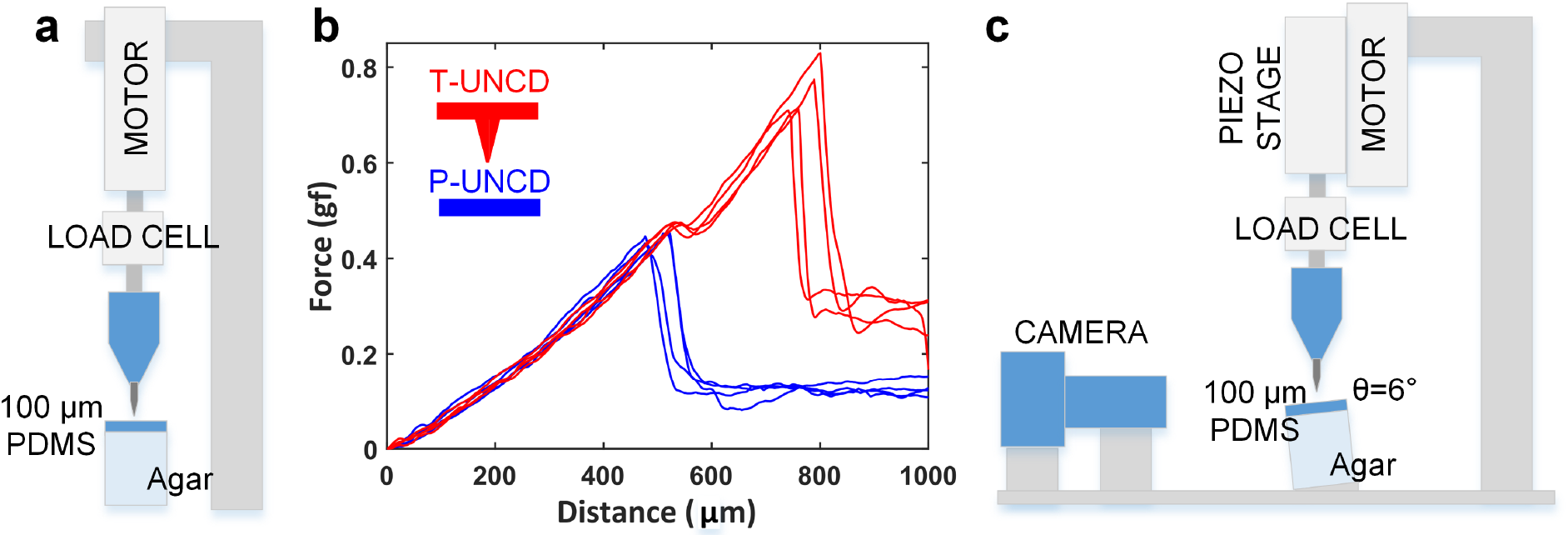
Measurement setup and force-distance curve for insertion through phantom. **a,** Measurement setup with a motor, a load cell and a tissue phantom with 100μm-thick PDMS. The phantom uses 100-μm thick layer of Sylgard 184 silicone (1-2 MPa^7^) over 0.7% agar to represent a membrane and soft nervous tissue. **b,** Force-distance curve for P-UNCD and T-UNCD. T-UNCD showed two bumps, with the earlier one indicating the insertion of the planar tip. Insertion speed 0.01mm/s and acceleration 0.1mm/s^2^. **c**, Measurement setup that includes a camera with 4X macro lens to measure maximum compression distance. Piezomotor from PiezoConcept, Inc designed for high-precision microscope stages was used here to enable uni-axial sinusoidal oscillations of up to 70-μm peak-to-peak.

**Supplementary Figure 4.**
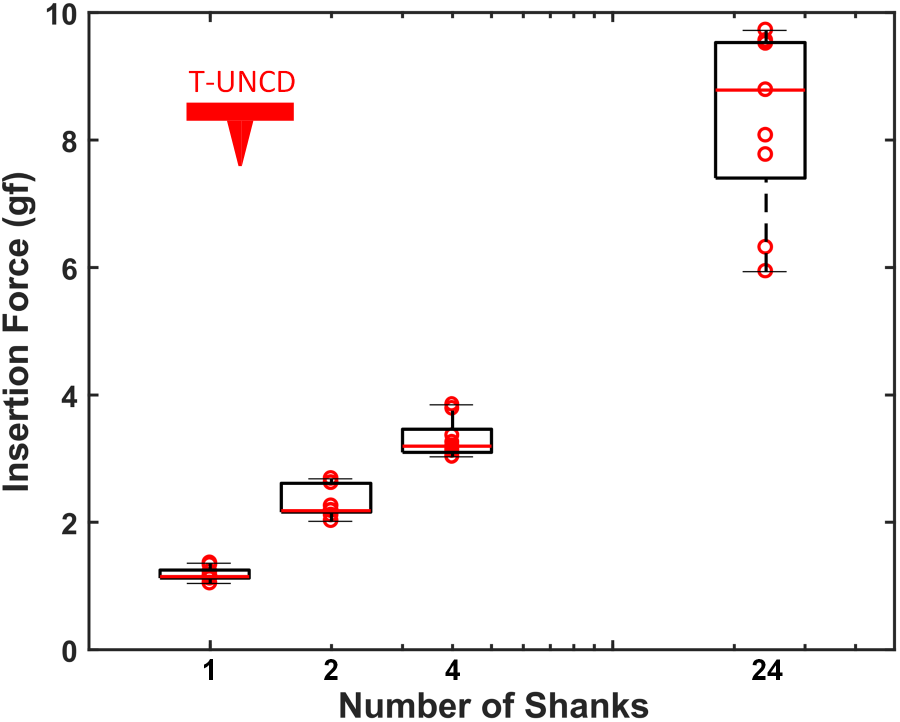
Phantom insertion force of 1,2,4 and 24 shanks for UNCD. To achieve 24 shanks, we arranged six 4-shank arrays in a hexagonal pattern. The force per shank in this case is 0.35 gf, a 66% and 72% force reduction relative to the 4-shank and 1-shank designs (Figure 3). This mechanism of local force reduction is yet unknown and may vary from shank to shank so more research on the local stress may help to reduce the size of a large number of shanks relative to the single shank version.

**Supplementary Figure 5.**
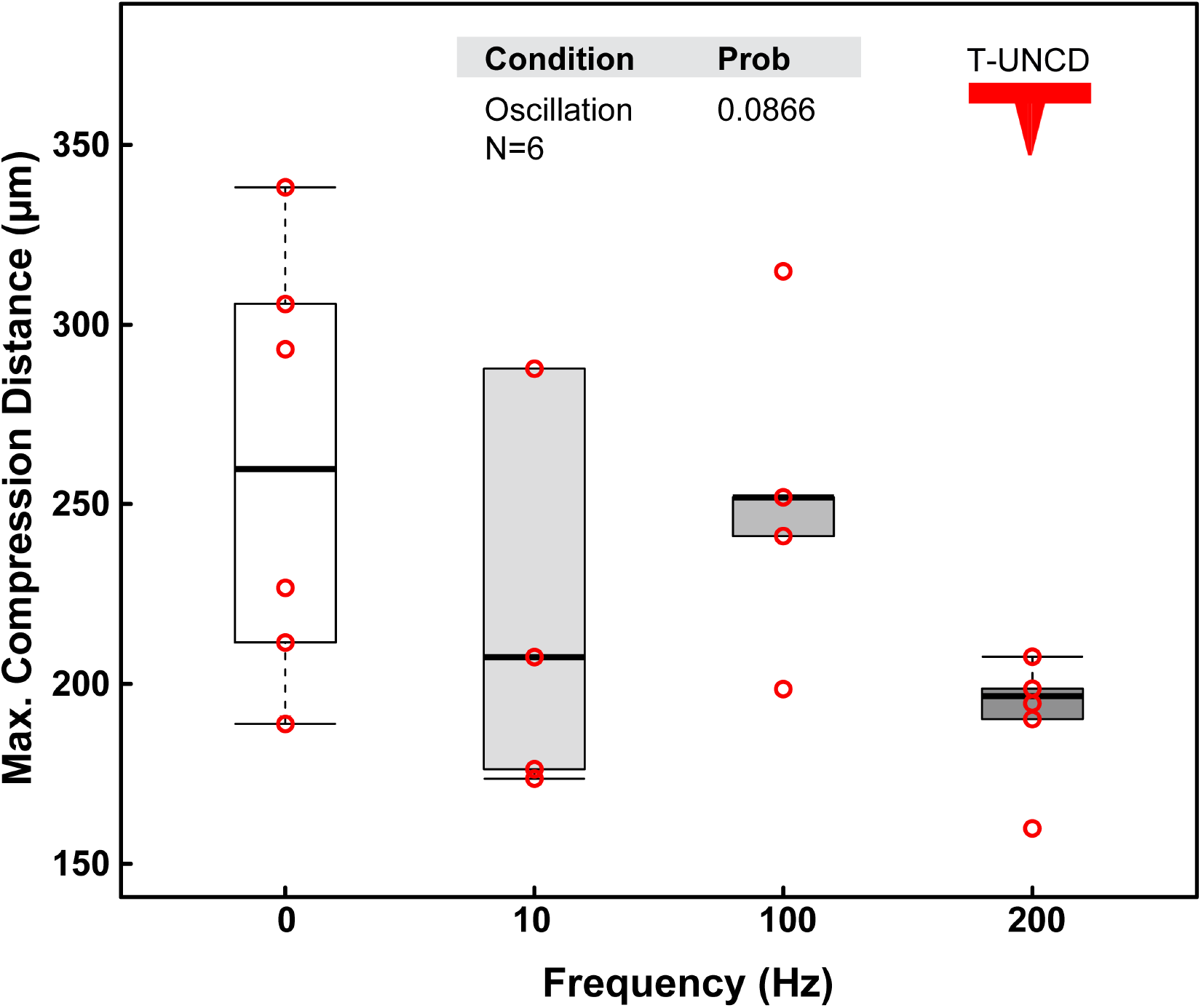
Maximum compression distance of UNCD shuttle with varying oscillation frequency at 2 mm/s in tissue phantom

**Supplementary Figure 6.**
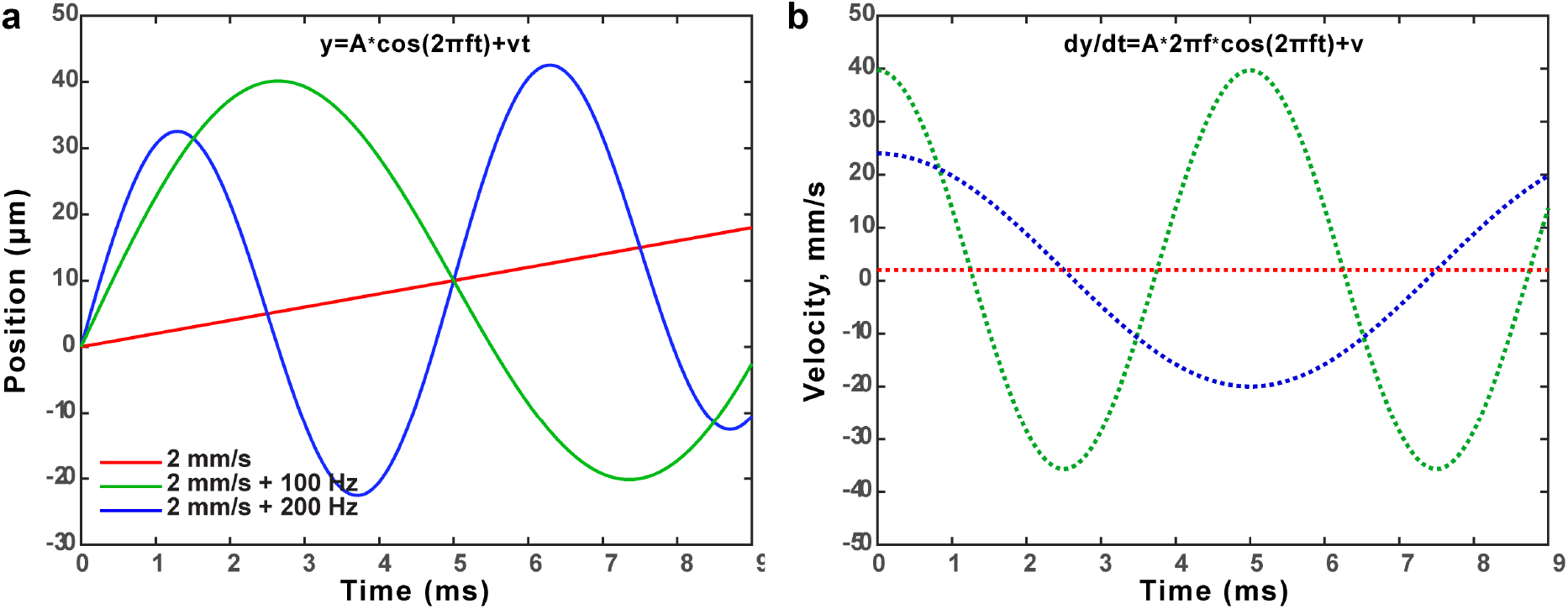
Kinematics of instantaneous insertion speed using a piezostage motor with uniaxial motion. The position equation and its first derivative are defined in the figures where A is maximum amplitude for a given oscillation. Beyond 200 Hz, the amplitude of this piezostage, HS1.70 from PiezoConcept, was dampened (see Supp. Table 1).

**Supplementary Table 1.** Peak Velocity and Acceleration of All Phantom Conditions Tested

|  | Frequency |  |  |  |
| --- | --- | --- | --- | --- |
|  | 10 | 100 | 200 | 400<sup>A</sup> |
| Max amplitude (μm) | 35 | 35 | 30 | 13.6 |
| Max speed (mm/s) | 2.2 | 22 | 38 | 34 |
|  | + linear sp | + linear sp | + linear sp | + linear sp |
| Max acceleration (m/s<sup>2</sup>) | 0.1 | 14 | 47 | 86 |

Note A: 400 Hz was only tested at 0.01 mm/s and only for 2 trials. Given it was a slower speed than 200 Hz and showed no improvement in those 2 trials we dropped this condition from the study.

**Supplementary Figure 7.**
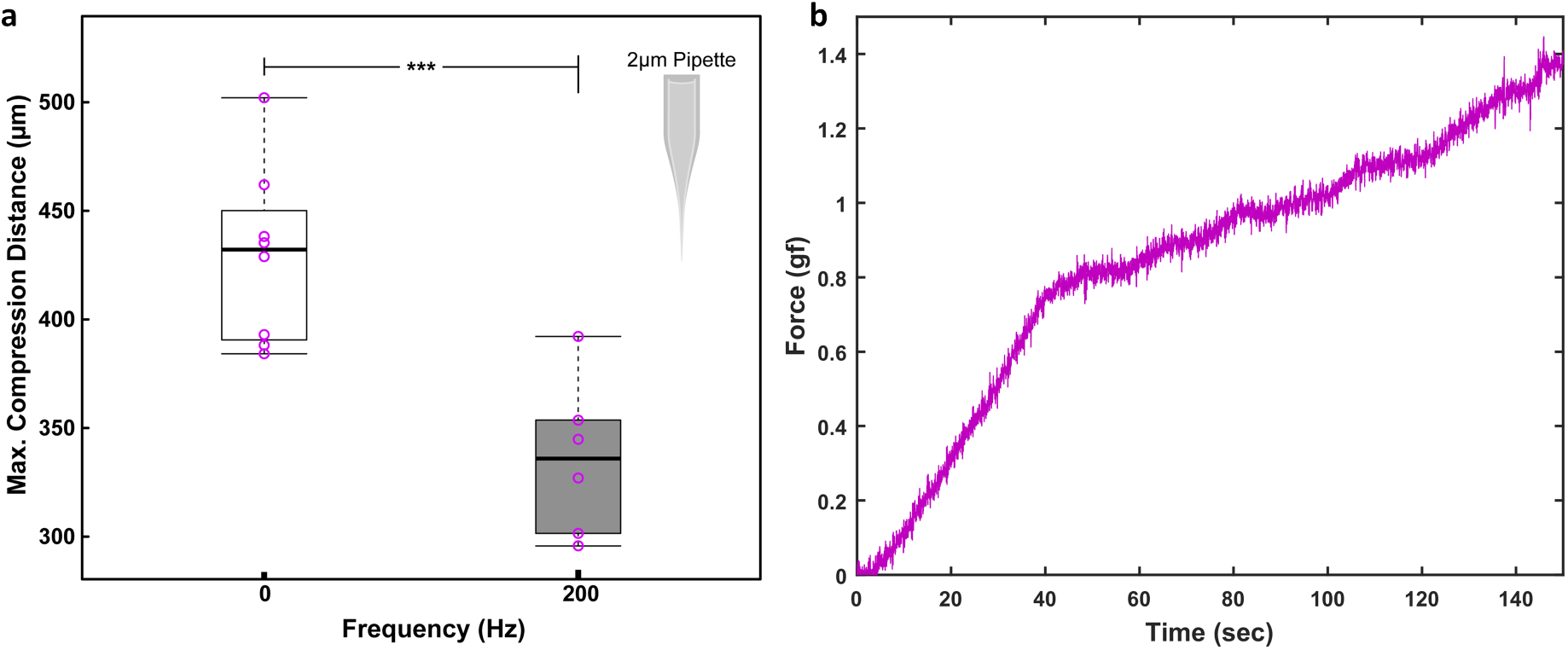
Oscillation and force testing of a 2-μm tip pipette at low speeds in tissue phantom. Glass pipettes are still a common implantable sensor for neuroscience and makes a useful model understanding the effect of device cross-sectional area, which is monotonically increasing. **a,** Oscillation significantly reduces the maximum compression distance (p=0.00074, N>6). **b,** While the tip is very sharp, we observe that the larger cross-sectional area dominates the force response, as reported by others^8^.

**Supplementary Figure 8.**
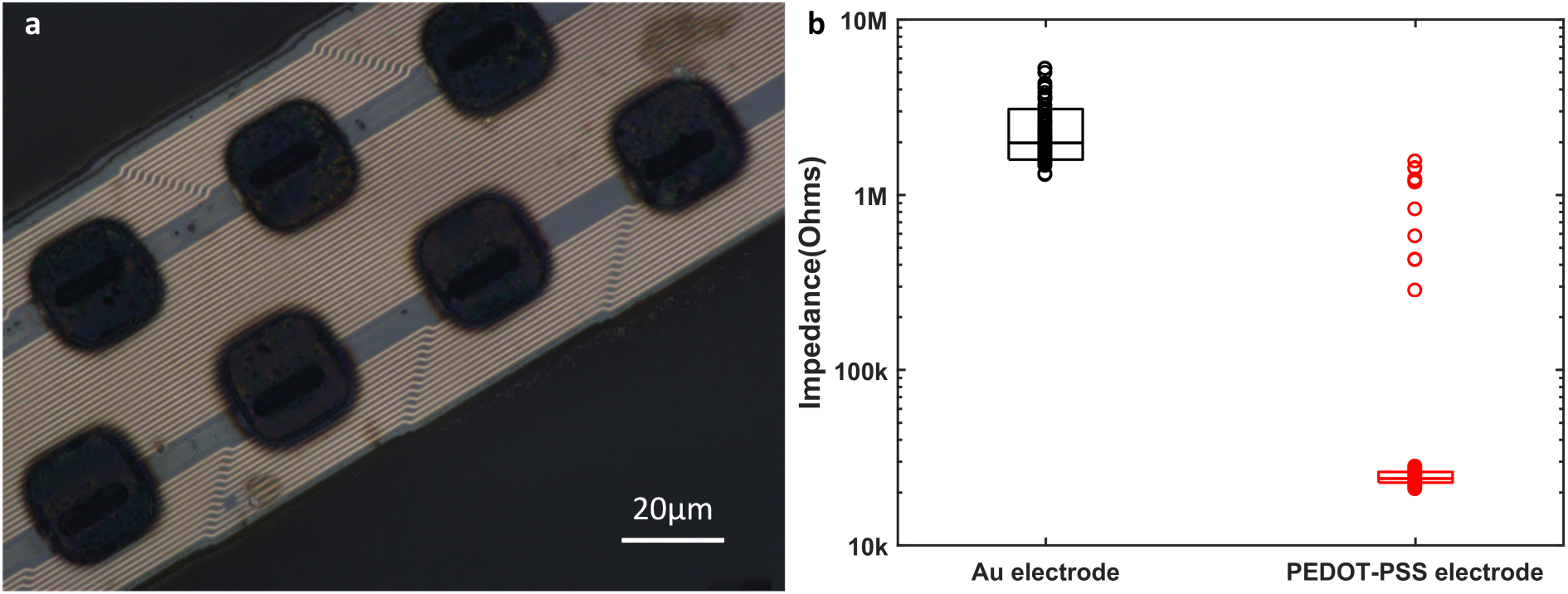
Picture and impedance of PEDOT:pTS coated electrodes. **a,** Poly(3,4-ethylenedioxythiophene: Sodium p-toluenesulfonate was plated on Au electrodes with a current of 690 nA and duration of 10 minutes as described in Patel, et. al 2016^9^. **b,** Impedances at 1kHz for non-coated Au electrodes and PEDOT-pTS coated electrodes, N=120. Top, middle and bottom lines of box plots indicates 90% value, harmonic mean, and 10% value, respectively. For the PEDOT-pTS coated electrodes, 14 electrodes with relatively high impedance were excluded as outliers.

### Supplementary Note 3: Detailed results of feline in-vivo experiments

This note includes additional descriptions of the electrode performance. In the first animal experiment a total of 28 different channels had a unit out of 42 functional channels during six sensory input trials. The per trial average was 11 units across 8 channels with single unit amplitudes between 35-323 μV peak-to-peak (mean 68 μVp-p, median 47 μVp-p).

In the second animal experiment an average of 78 units across 32 channels were observed during each of eight evoked trials during testing with the electrode in a fixed position, with single unit amplitudes between 39-1313 μV peak-to-peak (mean 123 μVp-p, median 80 μVp-p). Across multiple trials in this fixed position, a total of 48 different channels had a unit (167 non-unique units observed) out of 59 functional channels, spanning 1.06 mm in depth. When this same probe was retracted, units clearly moved between electrodes, including appearing on electrodes that were deeper in the tissue, suggesting that the tip of the array may have initially been in an inactive region of the DRG or in the ventral root (Fig. 7a). During movement of the probe, a total of 249 units were identified, with all 59 functional channels having activity across the full 1.2 mm span. Single units during the probe movement trial were within 50-1273 μV peak-to-peak (mean 199 μVp-p, median 125 μVp-p).

As the channels within the electrode array were separated by 40 μm, some units appear on multiple channels and we used careful manual sorting followed by comparison of waveform shapes across neighboring channels to report putative units. High-density arrays, like that used here, are best employed using an unbiased polytrode analysis as others have demonstrated^10–12^ to eliminate double counts but also to increase the number of unique identifiable units when performing an in-depth mapping study^13^.

**Supplementary Figure 9.**
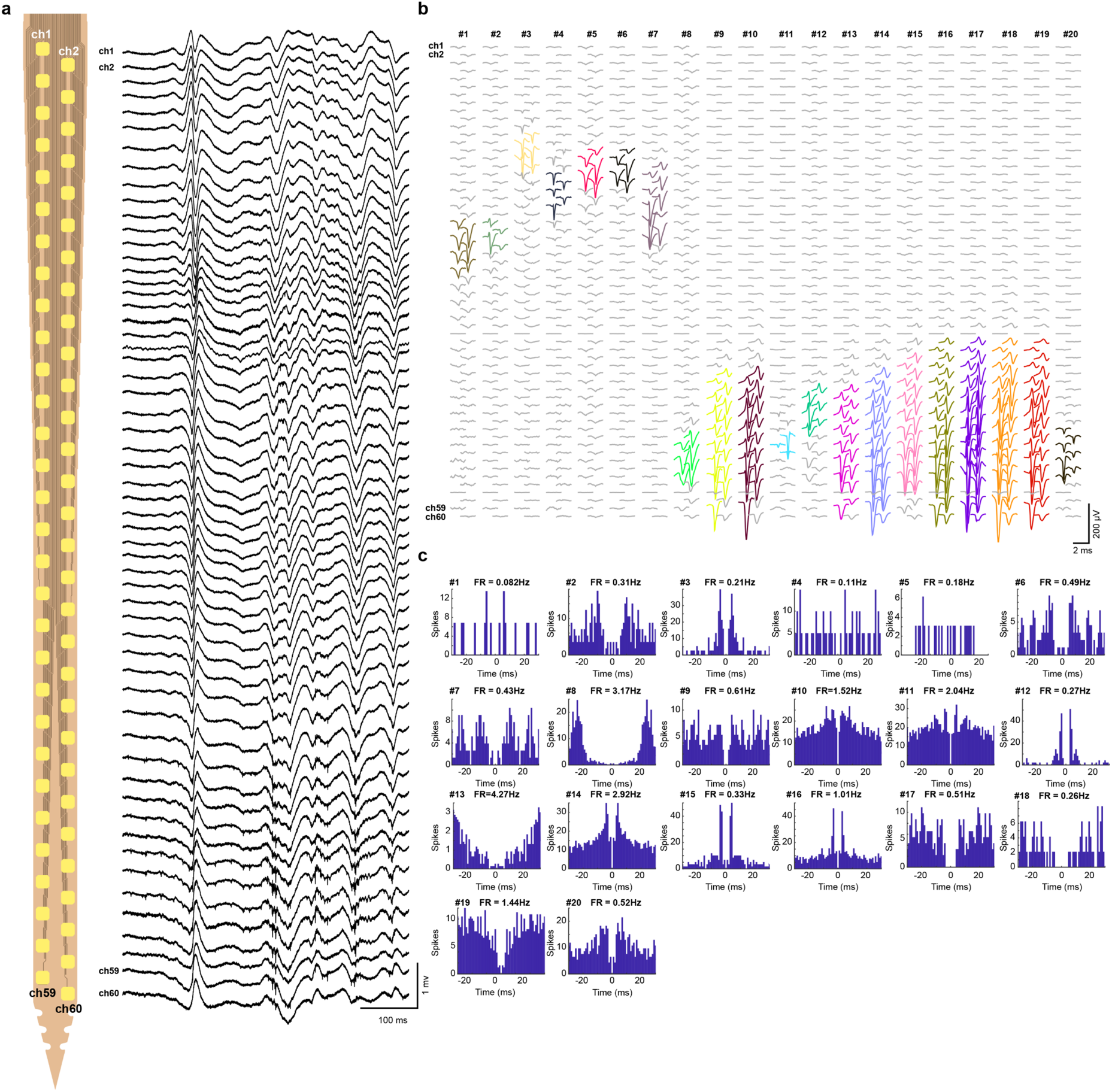
Recorded neural signal from somatosensory cortex of rat. **a,** Wide-band (0.3 – 3000 Hz) spiking activity recorded on a flexible electrode from S1FL region of an anesthetized rat (right) and a schematic of the probe sites (left). Note the spiking activity at the bottom of the shank. **b,** Recording of high number of well-isolated single units. The spiking data was quantified for 20 putative single units recorded simultaneously from S1FL (same animal and session as in **a**). Plots show average single-unit triggered waveforms (n = number of spikes, at least 500) on all channels. Note the single unit activity is present across multiple channels but the larger than expected range is due to the movement of probe relative to the brain tissue over the course of 30 minutes. **c,** Auto-correlation histograms of the 20 putative single units (same as **b**). Mean firing rate (FR) value is shown for each unit above the histogram.

**Supplementary Video 1 | Example of insertion failure due to tissue movement during *ex-vivo* trial.** The insertion speed was 0.01mm/sec and the travel distance was 1.5mm. The video showed the buckling failure of P-UNCD during the insertion trial into DRG L7 explanted from a feline. Link for video will be provided in a peer-reviewed journal article.

**Supplementary Video 2 | Side-by-side comparison of UNCD insertion without and with 200 Hz oscillation.** The linear speed of insertion was 0.01mm/s for both. Link for video will be provided in a peer-reviewed journal article.

**Supplementary Video 3 | In vivo insertion through rat dura and retraction leaving flexible array implanted.** The polyimide electrode array on T-UNCD shuttle was implanted into the motor cortex and the T-UNCD shuttle was successfully retracted, leaving the polyimide array inside. The insertion was conducted at 0.01mm/s speed with 200Hz oscillation. Retraction occurred in steps of 50 to 100 μm to ensure separation was visible before larger movements were made. Link for video will be provided in a peer-reviewed journal article.

